# Cell Fate Determining Molecular Switches and Signaling Pathway in Pax7-expressing Somitic Mesoderm

**DOI:** 10.1101/518084

**Authors:** Cheuk Wang Fung, Han Zhu, Shao Pu Zhou, Zhenguo Wu, Angela R. Wu

## Abstract

Pax7-expressing progenitor cells in the somitic mesoderm differentiate into multiple lineages, such as brown adipose tissue, dorsal dermis, as well as muscle in the dorsal trunk and the diaphragm; however, the key molecular switches that determine and control the process of lineage commitment and cell fate are unknown. To probe the mechanisms behind mesoderm development, Pax7^creER^/R26-stop-EYFP embryos were tamoxifen-induced at E9.5 to label Pax7+ cells for lineage tracing and collected at later time points for analysis. The YFP-labelled cells which belonged to the Pax7 lineage were enriched by fluorescence-activated cell sorting (FACS) and subject to single-cell RNA profiling. We observed that a subpopulation of cells differentiated into the myogenic lineage, showing *Myf5* expression as early as E12.5, whereas the rest of the population was fibroblast-like and appeared to be the early stage of the adipogenic and dermal lineages. Cells at E14.5 had distinct myogenic populations that expressed *Myod1* and *Myog;* we also identified other populations with *Ebf2* or *Twist2* expression, which could belong to adipogenic or dermal lineages, respectively. Cell surface markers were also found for each specific lineage, providing insights in sorting strategy for lineage-of-interest for further functional evaluation. Adipogenic lineage was successfully sorted with a combination of Pdgfra and Thy1 antibodies. In addition, we found that upregulation of Wnt signaling pathway activity is dynamically regulated in dermal lineage. Finally, transcription factors that could potentially drive, or reprogram cell fate, were identified at different developmental time points.

**Summary statement:** Investigation of Pax7 lineage transcriptomic profile at single-cell level identified multiple cell types, fate commitment time point, surface markers, transcription factors and signaling pathways that determine cell fate.

## Introduction

One longstanding and interesting question in developmental biology is how cell fates are chosen and specified from a common progenitor population. During embryogenesis, cell fate determination, cell proliferation, and cell survival are highly controlled processes in which Pax3 and Pax7, two highly similar Pax family transcription factors, are known to play important roles (Buckingham and Relaix, 2007). Pax3 expression begins in the presomitic mesoderm, while expression of Pax7 emerges later in the dermomyotome (Fan and Tessier-Lavigne, 1994; Lepper and Fan, 2010). At E10.5, a highly proliferative population of cells that express both Pax3/Pax7 can be found in the central part of dermomyotome, and these cells later contribute to the myogenic lineage, but lineage-specific markers such as Myf5 and Myod1 are not yet expressed at this early time point (Relaix et al., 2005). Other studies later demonstrated using CreER/LoxP lineage tracing that Pax7 expressing cells are capable of multiple cell fates, contributing to at least three lineages: interscapular brown fat, dorsal dermis and skeletal muscle (Lepper and Fan, 2010). Interestingly, cells expressing Pax7 at different developmental time points have different cell fate potential: Pax7-expressing cells early in development, at E9.5-E10.5, are able to contribute to dorsal dermis, BAT, as well as skeletal muscle; after E12.5, however, cells expressing Pax7 predominantly give rise to skeletal muscle (Lepper and Fan, 2010). This shift in cell fate potential suggests that different waves of Pax7-expressing cells emerge throughout development, and that each wave may be intrinsically different. In addition, Pax7 cells were also reported to give rise to olfactory neurons and neural crest cells (Murdoch et al., 2010; Murdoch et al., 2012; Relaix, 2004). These previous studies all confirm that the Pax7 lineage is able to give rise to various tissue types, and that fate commitment for each type happens in a time-dependent manner. However, the specific timing, as well as the detailed mechanisms driving multiple fate specification within the Pax7 lineage remain poorly understood. To elucidate these mechanisms requires knowledge of the molecular state of early lineage committed progenitor cells, and until now some of these progenitors have been challenging to identify and isolate for profiling and functional characterization.

Of the tissue types arising from the Pax7 lineage, the myogenic lineage is the most extensively studied. Pax3/7 expression appear to persist in the myogenic lineage, while cells adopting other fates quickly downregulate Pax3/7 expression (Buckingham and Relaix, 2007). The paradigm of Myf5 and Myod1 as the key fate determination factors, serving as upstream regulators of Myog and as markers of myogenic lineage commitment, is also well established (Buckingham and Relaix, 2007; Cossu et al., 1996). Linking these two observations, we recently demonstrated that Pax7 expression serves as a molecular switch between muscle precursors and brown adipocytes (An et al., 2017): Pax7 directly upregulates Myf5 and Myod1, which suppress brown fat fate choice by inhibiting Prdm16 through targeting E2f4 (An et al., 2017). This Pax7-Myf5/Myod1-E2f4-Prdm16 axis explains why Pax7 expression is maintained in muscle lineage, but not in brown adipose tissue (BAT) (Buckingham and Relaix, 2007). The development and differentiation of BAT depends heavily on networks of transcription factors including EBF2, PRDM16, PPARg and CEBPb (Wang and Seale, 2016), and intracellular signaling, such as BMP7 (Tseng et al., 2008); however, the early specification between BAT and muscle remains elusive. Our previous work demonstrated cell fate reprogramming between BAT and muscles (An et al., 2017), and we would like to further investigate in this direction. In contrast, much less is known about BAT and its progenitors. Recently, molecular signatures of preadipocytes that appear have been elucidated in mature fat depots, including the identification of specific cell surface markers and transcription factors of importance (Gupta et al., 2012; Rodeheffer et al., 2008). It is unknown, however, whether these signatures are also representative of embryonic preadipocytes, since even within the same lineage embryonic and adult tissue can greatly differ. One example is Sca1, which labels preadipocytes in mature fat tissue, but is not expressed in embryonic fat progenitors (Rodeheffer et al., 2008; Schulz et al., 2011; Wang et al., 2014). The transcription factor Ebf2 was identified as being important in brown fat development (Rajakumari et al., 2013), and was also reported as an early marker for embryonic brown fat progenitors since Ebf2+ cells isolated from dorsal anterior regions display strong adipogenic potential (Wang et al., 2014). However, Ebf2 was also found to be involved in embryonic neurogenesis (Chuang et al., 2011; Jimenez et al., 2007). Thus, it is likely that additional transcriptional mechanisms exist to define neural and brown fat progenitor identities within the Pax7 lineage. Like BAT, dermal tissue is of fibroblastic lineage and can arise from the Pax7 lineage. To date, the most well established dermal marker is Twist2, a bHLH transcription factor downstream of the Wnt/beta-catenin pathway (Budnick et al., 2016; Li et al., 1995; Olivera-Martinez et al., 2004). The critical role of the canonical Wnt/beta-catenin pathway in dermal development has been examined extensively, and its importance in ventral, cranial and dorsal dermal cell development has been demonstrated (Atit et al., 2006; Ohtola et al., 2008; Tran et al., 2010). Embryonic dermal progenitors are marked by expression of En1 and Twist2, both of which are nuclear factors that cannot be used for cell isolation. One surface marker that labels the embryonic dermal lineage is Pdgfra, which is a receptor that is expressed in many cells of the fibroblastic lineage including preadipocytes. Thus, it remains challenging to directly isolate and investigate the lineage-specific progenitors of Pax7 origin (Budnick et al., 2016).

With the advent of single cell RNA sequencing technology (scRNA-seq), it is possible to obtain transcriptome signatures at single cell resolution and unravel the molecular heterogeneity within a mixed cell group without having to first sort them based on surface marker expression. Cells with similar gene expression can be grouped based on their transcriptomic signatures in an unbiased fashion, allowing identification of novel cellular subpopulations, sometimes even leading to discovery of surface markers that enable subsequent purification and functional characterization of these cells-of-interest. This is particularly useful for studying cells-of-interest when little is known about them or when they are a minor proportion of the total heterogeneous population, such as the aforementioned lineage-specific embryonic progenitors of Pax7 origin. In this study, we employed scRNA-seq combined with lineage tracing to perform transcriptional profiling of the Pax7 lineage throughout embryonic development. We specifically focused on the Pax7-expressing progenitors at early embryonic time point (E9.5) and examined their progenies at different developmental stages (E10.5, E12.5, E14.5). We identified four types of cells with distinct cell fates and reconstructed their lineage progression trajectories from a common progenitor group. For each cell population, we discovered unique cell markers to classify them, enabling new purification strategies for early lineage-specified progenitor cells. Importantly, from the transcriptomic data, we elucidate important transcriptional programs and signaling pathways that drive cell fate decision at early stage, expanding our understanding of fate commitment mechanisms in these lineages during early development.

## Results

Using a Pax7^creER^/R26-stop-EYFP mouse, wherein tamoxifen-induced YFP expression labels those progeny cells descended from Pax7-positive progenitors, we induced YFP expression at day E9.5 and performed scRNA-seq profiling on Pax7 progeny at E10.5, E12.5, and E14.5. We profiled a total of 728 single cells using Smartseq2, which provides full-length transcripts, and a total of 3140 single cells using the 10X Genomics platform, which provides higher throughput. Using unsupervised clustering, we determined 8 distinct cell clusters (Fig. 1A-D). The cell type identify of each cluster was assigned based on their unique gene expression pattern, using either established marker genes, Gene Ontology analysis, or other previously reported findings (Fig. 1E-T). The cell clusters comprise four different tissue lineages, all of which have been previously reported to arise from the Pax7-lineage: myogenic, dermal, adipogenic, and neuronal. Almost all cells from E10.5 embryos were attributed to Pax7 progenitor cells and expressed no lineage-specific genes. Due to the small size of the embryo and the low numbers of cells available for analysis at this time point, we were not able to observe any intrinsic differences within the Pax7 progenitor cell population in our data. We subsequently used likelihood-ratio test for differential gene expression analysis (McDavid et al., 2013) to find novel gene markers, cell surface markers, and transcription factors that may be used to isolate specific lineages, or for lineage reprogramming for the other cellular subpopulations. We also not only confirmed that Wnt signaling is crucial for dermal lineage identity, but also found evidence that Wnt pathway associated factors are important in the fate commitment decision defining the dermal lineage from the adipogenic lineage at early embryonic development.

**Fig. 1.**
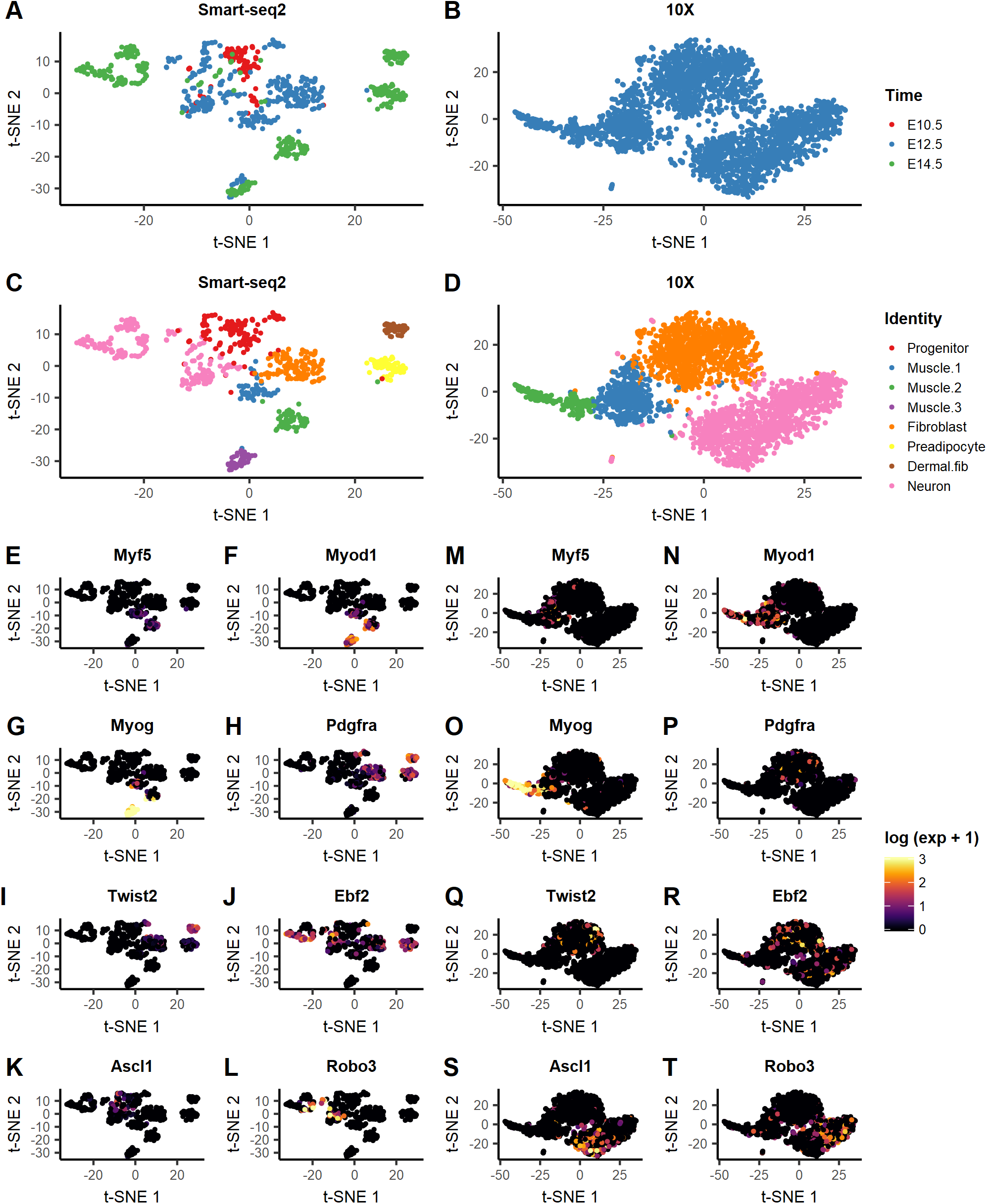
Transcriptomic profile of Pax7 lineage at single-cell level. (A-D) Cells from Pax7 lineage across E10.5, E12.5, and E14.5 labeled by their collection time point after t-distributed stochastic neighbor embedding (t-SNE) visualization in Smart-seq2 and 10X data. (A) Smart-seq2 data contained cells from all three time points, while (B) 10X data consisted cells from E12.5 only. (C-D) Cells labeled by their identity after referencing gene markers from previous findings and Gene Ontology analysis to clustering result in both Smart-seq2 and 10X data. (E-L) Gene markers to identify cell lineages in Smart-seq2 data, and (M-T) in 10X data.

### *Ascl1* and *Robo3* pinpoint the neuronal lineage

Previous studies have suggested that the Pax7 lineage contributes to olfactory neurons and neural crest cells. Our data revealed multiple cell clusters with expression of the proneuronal gene *Ascl1* and the axon guidance receptor gene *Robo3* (Fig. 1K-L,S-T), supporting previously observations of neuronal lineage contributions from Pax7 progenitors (Murdoch et al., 2010; Murdoch et al., 2012; Relaix, 2004). Gene Ontology enrichment analysis also supported the neuronal identity of these cell clusters (Fig. S1).

These neuronal-lineage cells represent a surprisingly large proportion of total cells profiled. Among them we observed a large number of Ebf2-expressing cells (over 20% of all cells classified as neuronal are Ebf2+), which were previously found to be crucial for olfactory neuron development: Ebf2 knockout mice have defective olfactory neurons that are unable to project to the dorsal olfactory bulb (Wang, 2004). This supports the view that the Pax7 lineage also gives rise to olfactory epithelial (OE) progenitors. An earlier study of Pax7-expressing cells and their specific contribution to the post-natal OE observed that Pax7-positive cells give rise to fewer postnatal OE cells compared to Pax7-negative cells, but the authors also note that this could be due to a depletion in the reservoir of Pax7 progenitors with age (Murdoch et al., 2010). Although it is possible that the single cell tissue dissociation procedure introduced cell-type specific bias that led to over-representation of these cells in the dataset, the abundance of Ebf2-positive neuronal progeny from Pax7 progenitors in our data is intriguing. Furthermore, the number of Ebf2-positive neuronal cells appears to increase from E12.5 to E14.5 (48.9% at E12.5 to 66.9% at 14.5). Given these observations, the degree to which the Pax7-positive lineage contributes to OE early in development warrants additional investigation. The remaining neuronal cells that are Ebf2-negative are likely to be Ascl1- and Crabp1-expressing uncommitted neural progenitors, or Robo3-expressing neural crest cells (Fig. 1J, R, L, T).

### Subpopulations in the myogenic lineage are defined by differential expression of *Myf5, Myod1*, and *Myog* across developmental time

*Myf5* and *Myod1* gene are well-studied transcription factors that determine myogenic cell fate by regulating *Myog* expression (Buckingham and Relaix, 2007; Cossu et al., 1996; Kablar et al., 2003). In our data, *Myf5* and *Myod1* were found upregulated exclusively in three specific cell clusters spanning from E12.5 to E14.5 in Smartseq2 data, and in two clusters in 10X data with only cells from E12.5 embryos (Fig1. E-G, MO). Differential gene expression analysis of these cell clusters relative to other identified cell clusters produced Gene Ontology terms related to muscle structure and tissue development, and muscle cell differentiation with high significance (Fig. S2), which further confirms their myogenic identity. Their respective expression levels of *Myf5* and *Myod1* indicated that they are early (Muscle.1), intermediate (Muscle.2), and late (Muscle.3) myogenic cell subpopulations (Fig. 2A-B), corresponding closely with the embryonic time at which the cells were sampled. Early myogenic progenitors that are only found at E12.5 expressed lower levels of Myf5 and Myod1 as compared to the intermediate group, which is only found at E14.5. In contrast, the late or more well-differentiated myogenic subpopulation that can be found at both E12.5 and E14.5 loses expression of Myf5 but upregulate the expression of Myod1 and Myog (Fig. 1E-G, M-O). This observed upregulation of Myod1 and Myog over time is consistent with the classical understanding of myogenic lineage commitment and differentiation (Buckingham and Relaix, 2007; Cossu et al., 1996; Kablar et al., 2003).

**Fig. 2.**
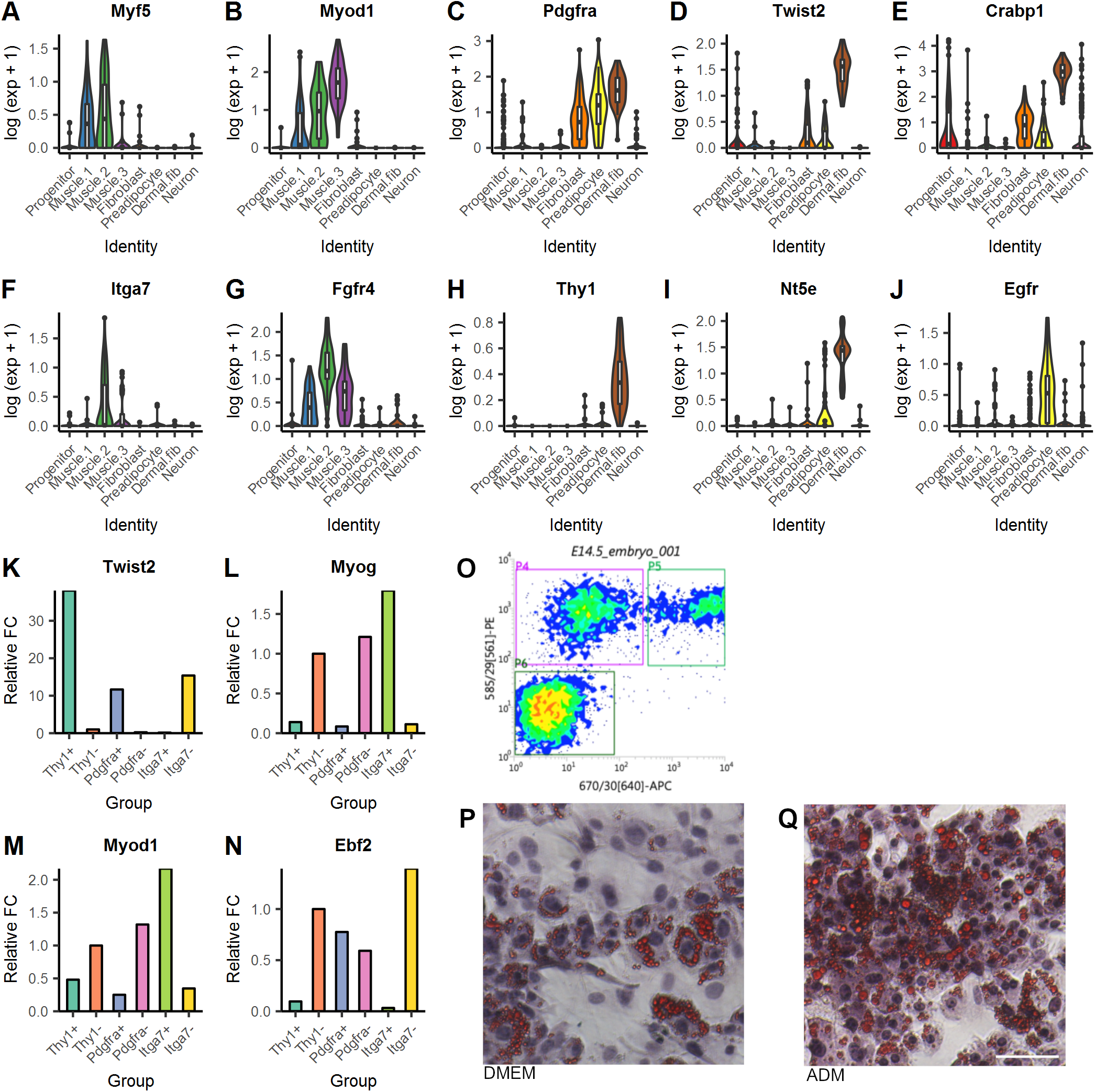
Gene expression level and cell surface markers for sorting lineages. (A-J) Expression level of genes differentially expressed in Smart-seq2 data. (A-B) Expression of *Myf5* and *Myod1* defined early (Muscle.1), intermediate (Muscle.2), and late (Muscle.3) myogenic lineage. (C) *Pdgfra* was expressed in fibrboblasts, dermal and adipogenic lineage cells. (D-E) *Twist2* and *Crabp1* were co-expressed in dermal lineage cells. (F) *Itga7* which reported to mark skeletal myoblasts (Blanco-Bose et al., 2001) was surprisingly expressed only in one cluster in myogenic lineage. (G) *Fgfr4* was found upregulated in all three clusters in myogenic lineage, showing potential to replace *Itga7* to label myogenic lineage. (H-I) *Thy1* and *Nt5e* were found exclusively upregulated in dermal lineage cluster. (J) *Egfr* was expressed only in adipogenic lineage cluster. (K-N) Real-time PCR result on sorted cells, surface markers used was on x-axis. Thy1^+/-^ were cells first Pdgfra+ enriched. Fold change was normalized to Thy1-expression on tested qPCR primers. (L-M) *Myog* and *Myod1* were upregulated only in Itga7+ cells. (N) *Ebf2* had higher expression in Thy1- and Itga7-cells. *Ebf2* expression from Itga7-cells might come from neuronal lineage. (O) Density plot of Pdgfra/Thy1 double sort. Pdgfra antibody reporter signal was reflected on y-axis, while Thy1 antibody reporter signaling was reflected on x-axis. (P-Q) Oil Red O staining of Pdgfra+/Thy1-cell culture. (P) Oil droplet staining in cells cultured in DMEM medium and (Q) in adipogenic medium (ADM) previously described (An et al., 2017). Scale bar: 50 μm in Q. P and Q share the same scale.

Furthermore, the early myogenic cluster is the only non-neuronal lineage at E12.5 that is lineage-committed at this early time point, as indicated by Myf5 expression (Fig. 1E, M, Fig. 2A); the remaining cells profiled at E12.5 do not express any distinctive markers for either the dermal or adipogenic lineages. Furthermore, at E12.5 there are already some Myog-expressing late myogenic cells, suggesting that myogenic fate decisions occur between E10.5 and E12.5, and that myogenic cells diverge from the fibroblastic lineage as early as E12.5, but lineage specification within the fibroblasts into either dermal or adipogenic lineages occurs later. One particularly provocative observation is that some cells in the late myogenic group (Muscle.3) are found at E12.5 whereas the intermediate group (Muscle.2) is only found at E14.5. Given that previous studies have shown Myf5-expressing cells can give rise to both skeletal muscle and BAT (Seale et al., 2008), but our data shows the BAT lineage commitment occurring later than the myogenic lineage, it could indicate multiple waves of myogenic lineage commitment via intermediate cellular subpopulations with different fate potential. A more detailed explanation is provided in the discussion section.

### Fibroblast progenitors marked by Pdgfra share common expression signatures with preadipocytes and dermal progenitors

A large cluster of cells from E12.5 expressed high levels of Pdgfra, a marker of fibroblasts (Fig. 1H, P, Fig. 2C). We also found sparse expression of both *Twist2* and *Ebf2* in this population (Fig. 1I-J, Q-R); this suggests transcriptional priming for later lineage specification toward either dermal or adipogenic lineage given the prominent *Twist2* and *Ebf2* expression respectively in these lineages.

### Dermal and adipogenic lineages are differentiated by *Twist2/Crabp1, Ebf2* and *Thy1* expression

*Twist2* (Dermo-1) is a transcription factor previously reported as a marker for dermal differentiation (Budnick et al., 2016; Li et al., 1995; Olivera-Martinez et al., 2004). Its importance in dermal development has been demonstrated in *Twist2-/-* mice with atrophic dermis (Šošić et al., 2003). In a subpopulation of cells found at E14.5, *Twist2* expression is strongly expressed (Fig. 1I, Q), which suggests their involvement in dermal lineage. *Crabp1* expression coincides with *Twist2* expression in this subpopulation; CRABP1 protein is known to be expressed in embryonic dermis, and previously the earliest observable CRABP1 expression using immunostaining was at E13.5 (Collins and Watt, 2008). With scRNA-seq data, *Crabp1* gene expression is revealed in a subset of Pdgfra-positive fibroblast progenitors as early as E12.5 (Fig. 1H, Fig. 2C, E), highlighting the resolution advantages of this technique. Transcription factor *Ebf2* was reported as a marker for brown adipocyte precursor cells, and its function in establishing brown adipose precursor gene program was shown previously (Rajakumari et al., 2013; Schulz et al., 2011; Wang et al., 2014). Indeed, we observed strong Pdgfra/Ebf2 co-expression in one of the E14.5 cellular subpopulations that is Twist2-negative (Fig. 1H-J). Despite strong evidence of dermal and adipogenic lineage identity from literature, Gene Ontology enrichment analysis on the transcriptomic signatures of these cell clusters returned no terms related to either of the lineages (Fig. S3); we also did not observe expression of intermediate and mature brown adipocyte markers *Pparγ* and *Ucp1* in the Ebf2-positive cluster (Cristancho and Lazar, 2011; Kajimura et al., 2015; Wang and Seale, 2016). Evaluating the evidence collectively, we hypothesized that these two subpopulations identified at E14.5 are not mature dermis or BAT, but rather dermal and adipogenic progenitors.

Reliable cell surface markers are invaluable for any downstream analysis that involves sorting lineage-specific cells. To be able to isolate these Pax7-derived lineage-specified progenitors for validation, we performed differential gene expression analysis on the transcriptome of each cell cluster, and discovered differentially expressed surface marker genes to distinguish the myogenic, dermal, and adipogenic lineages. *Pdgfra* was found differentially expressed in fibroblast-like cells at E12.5 (Fig. 2C), dermal, and adipogenic lineages at E14.5, while *Fgfr4* was differentially expressed in all three myogenic lineage clusters (Fig. 2G). Between dermal and adipogenic lineages, *Thy1* and *Nt5e* particularly defined the dermal lineage cluster (Fig. 2H-I), while *Egfr* defined the adipogenic lineage cluster (Fig. 2J).

To further investigate the feasibility of these factors as surface markers for cell purification, we sorted E14.5 embryos based on the hypotheses formed using scRNA-seq data: dermal lineage cells are Pdgfra+/Thy1+ while adipogenic lineage cells are Pdgfra+/Thy1-, and myogenic lineage cells are Itga7+ (a known marker of skeletal myoblasts (Blanco-Bose et al., 2001)). Real-time PCR on sorted populations showed *Myod1* and *Myog* were both highly expressed in Itga7+ cells and downregulated in Pdgfra+ cells (Fig. 2L-M), while *Twist2* and *Ebf2* were upregulated in Thy1+ and Thy1^-^ cells respectively (Fig. 2K, N). These results are consistent with our expectations.

To functionally validate the identity of the Pdgfra+/Thy1-sorted cell population, which we hypothesized to be the pre-adipogenic progenitors, we cultured these sorted cells *in vitro* in DMEM (Fig. 2O-Q). Upon reaching confluency, many of these Thy1-cells developed lipid-filled structures, as visualized by Oil Red O staining (Fig. 2P). We continued culturing a subset of these confluent Thy1-cells, swapping out DMEM for adipogenic medium (ADM) for two more days after confluency, and we observed drastically more lipid droplet-containing cells after culturing in adipogenic medium (Fig. 2Q). These results show both the spontaneous derivation of adipocytes from Pdgfra+/Thy1-cells in DMEM, as well as the high efficiency with which ADM culture can induce adipogenesis in these cells, serving as strong evidence that we have identified and found a sorting strategy for pre-adipogenic progenitors of the Pax7 lineage.

### Wnt/β-catenin activity is dynamically regulated in the dermal lineage

*Lef1* and *Axin2*, two important genes in Wnt signaling pathway, as well as Wnt inhibitors *Dkk1* and *Sfrp5*, were found upregulated in Pdgfra+/Thy1+ cluster at E14.5 that we hypothesized are the dermal fibroblasts (Fig. 3A-D). Both the dermal fibroblasts and preadipocytes appear to branch out from the earlier E12.5 fibroblast population, and given the known importance of canonical Wnt signaling in postnatal and adult dermal tissue, we postulate that Wnt signaling is the key determinant of cell fate for early fibroblastic progenitors facing a dermal fibroblast or preadipocyte fate choice. In postnatal mice, Lef1 mutants exhibit upregulation of Crabp1 gene expression, but in the presence of Lef1 expression, Crabp1 expression is lowered but not completely inhibited (Collins and Watt, 2008). Notably in our data, Lef1 expression is exclusive to this Twist2/Crabp1 double-positive group (Fig. 3A, Fig. 2D-E), supporting Wnt signaling as a dynamic and tunable mechanism for maintaining dermis tissue identity.

**Fig. 3.**
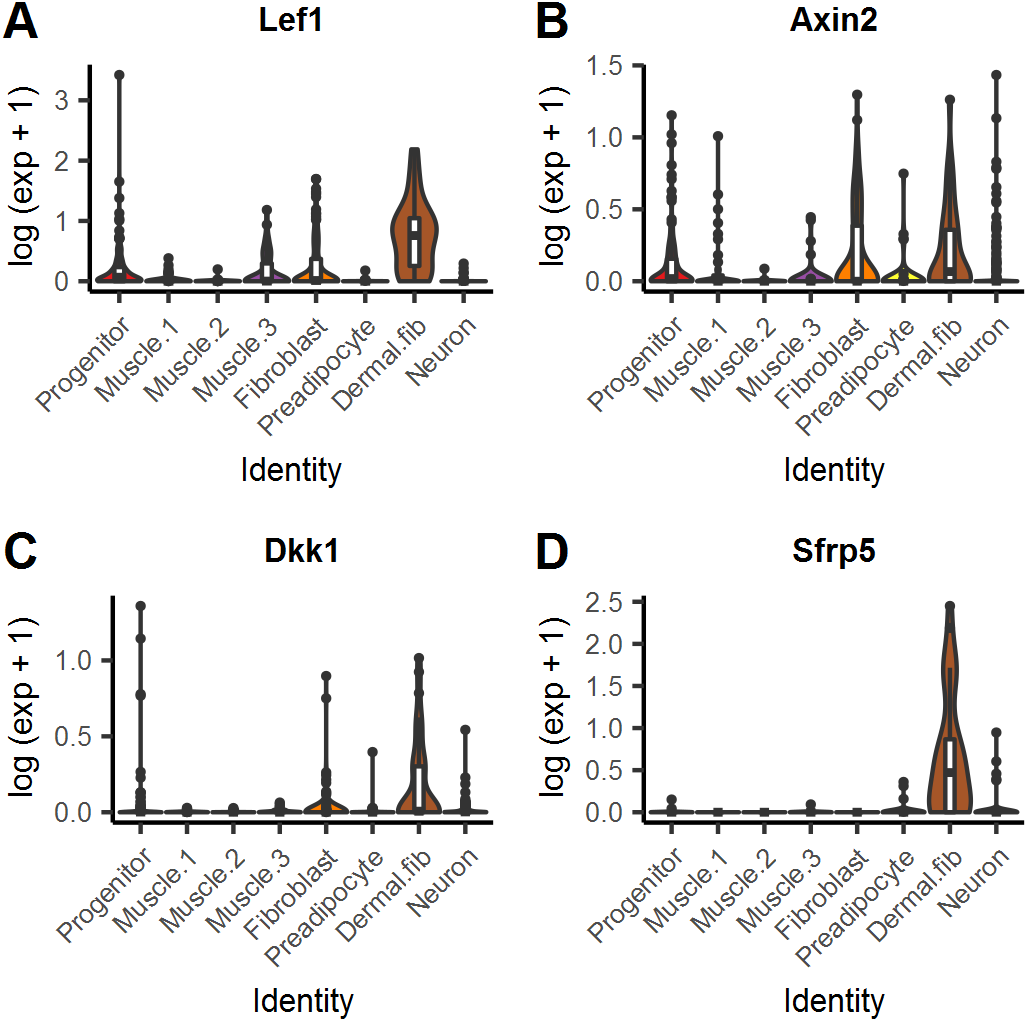
Wnt signaling pathway activity in dermal lineage. (A-D) Expression of genes involved in Wnt signaling pathway. (A-B) *Lef1* and *Axin2* are two important genes in Wnt signal transduction and is differentially expressed in dermal lineage. (C-D) *Dkk1* and *Sfrp5* are Wnt signaling inhibitors reported to regulate Wnt signaling activity dermal growth (Andl et al., 2002; Satoh et al., 2008), and were differentially expressed as well.

### Differential gene expression revealed key transcription factors for cell fate determination

Next, we investigated the transcription factors that were specific to the lineages, to yield mechanistic insight on the process of cell fate determination. Differentially expressed transcription factors were identified by referencing mouse transcription factor list obtained from TcoF-DB (Schmeier et al., 2017) and differentially expressed genes in each cluster. Since myogenic lineage become committed at E12.5, whereas dermal and adipogenic lineages committed at E14.5, transcription factors that are differentially expressed at E12.5 and E14.5 at the separate branch points would potentially point to transcription factors that drive the cell fate.

We found that at E12.5, when the myogenic lineage and fibroblast-like cluster separate, *Pitx3* was upregulated in early myogenic progenitor cells (Fig. 4B), whereas *Nfatc4* was upregulated in the early-fibroblast progenitor cluster (Fig. 4A). *Nfatc4* belongs to the family nuclear factor of activated T cells (NFAT) controlled by calcineurin, and have been shown to control cell fate of different tissue types, including T cells (Horsley and Pavlath, 2002; Serfling et al., 2006). It would not be surprising for *Nfatc4* to be the driver for fibroblast-like cluster at E12.5. *Pitx3* has been investigated for its importance in myogenesis at early embryonic stage, and deletion of *Pitx2* and *Pitx3* results in perturbed muscle development (L’honoré et al., 2014; L’Honoré et al., 2007). At E14.5, *Bcl11b* was enriched in dermal cluster, while *Meox2* was enriched in preadipocyte cluster. *Bcl11b* has been reported as a lineage commitment factor in T-cell development, while *Meox2* has been investigated on its angiogenesis-related function in brown adipocytes (Timmons et al., 2007). The differentially expressed transcription factors we have identified in each lineage may also play major roles in cell fate decision (Table 1). Our findings open up many new avenues of future investigation.

**Fig. 4.**
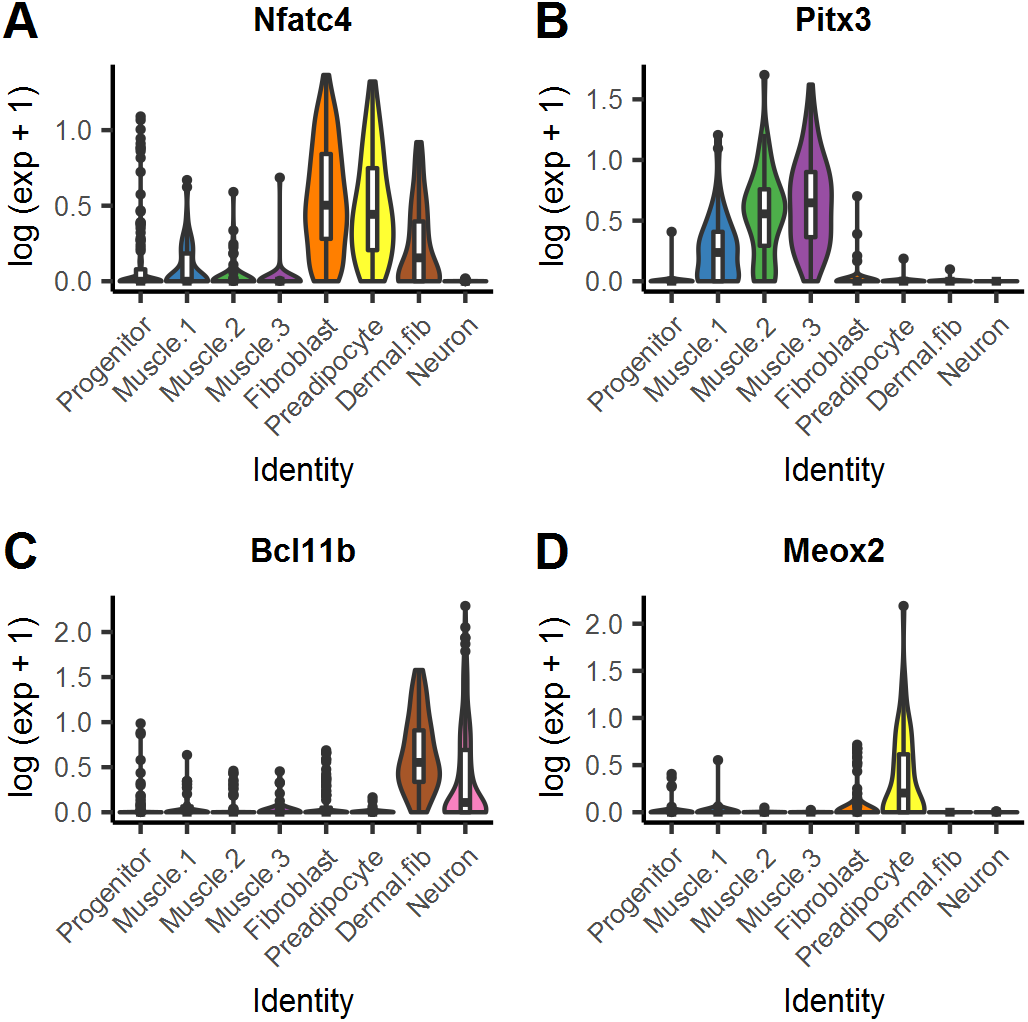
Key transcription factors that determine cell fate. (A-D) Gene expression of transcription factors differentially expressed in clusters from cells at (A-B) E12.5 and (CD) E14.5. (A) *Nfatc4* was differentially expressed in fibroblast-like cells at E12.5, and expression continued to dermal and adipogenic lineages to E14.5. (B) *Pitx3* was opposite to *Nfatc4* expression starting from early muscle cluster at E12.5, and continued expression in myogenic lineage at E14.5. (C) *Bcl11b* was found differentially expressed in dermal lineage cluster, while (D) *Meox2* was differentially expressed in adipogenic lineage cluster.

**Table 1.**
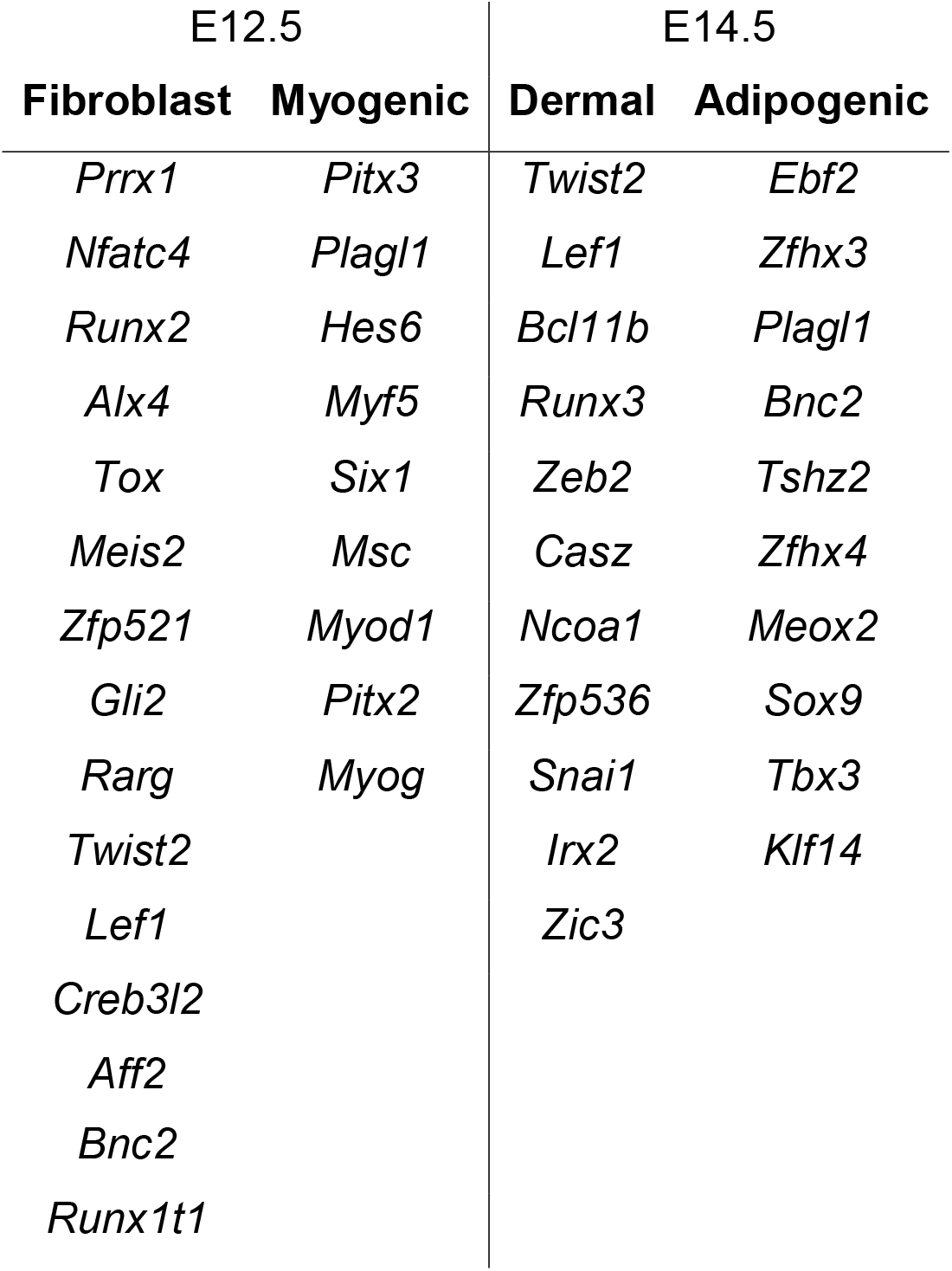
Differentially expressed transcription factors at different commitment time point.

## Discussion

Previous Pax7 lineage studies have highlighted the multipotency of Pax7-positive progenitor cells, which can develop into skeletal muscle, interscapular brown fat, and dorsal dermis (Lepper and Fan, 2010), and possibly olfactory neurons and neural crest cells (Murdoch et al., 2010; Murdoch et al., 2012; Relaix, 2004). Despite well-known cell fate commitment in Pax7 lineage, the time points when cells commit to a particular lineage are still inconclusive. By tracing and probing Pax7 lineage at multiple development time points in single-cell resolution, we here show the transcriptomic profile across Pax7 lineage development. Referencing previously studied gene markers identified the myogenic, dermal, adipogenic, and neuronal lineages, with discovery of lineage commitment at certain time points during embryonic development. Focusing on the three lineages, we found that lineage commitment Pax7+ progenitors happens between E10.5 and E14.5. During this time course, myogenic lineage branches out at E12.5 with *Myf5* upregulation, while the remaining cells separate at E14.5 and commit into either dermal or adipogenic lineage with *Twist2* or *Ebf2* upregulation. Our analysis agrees with previous findings on Pax7 lineage, and it formally shows for the first time the temporal relation between each lineage commitment. Intriguingly, we noticed that some cells in the more mature myogenic group (Muscle.3) are found at an earlier time point, E12.5, whereas the less mature myogenic group (Muscle.2) is found exclusively at a later time point, E14.5. Assuming that all Myog+ cells must be derived from a population of Myf5+/Myod1+ cells, then the Myog+ cells found at E12.5 are likely to be the result of an earlier wave of differentiation from Myf5+ progenitors arising between E10.5 and E12.5. The Myf5+ cells found at E12.5 (Muscle.1) can continue to give rise to a second wave of Myog+ cells, captured at E14.5. Previously, Myf5-expressing cells were shown to give rise to both skeletal muscle and BAT (Seale et al., 2008), but in our data, the adipogenic lineage commitment occurs much later than the myogenic lineage. The clustering of Myf5+ progenitors that are found at E14.5 (Muscle.2) as a separate group from those found at E12.5 (Muscle.1) indicates they are transcriptionally distinct; we confirmed that the difference is not due to cell cycle effects or batch effects. It is possible that the Myf5+ progenitors found at E14.5 are able to still switch to an adipogenic fate. Further investigation of Myf5 lineages is needed to explore this possibility.

In Myf5 lineage, previous studies demonstrated that Ebf2 protein is expressed in E11.5 embryos and Ebf2+ population dramatically expanded at E12.5 (Wang et al., 2014). Suggesting that brown adipogenic cell fate is determined by E12.5; however, our Pax7 lineage data showed low expression of *Ebf2* in non-neuronal cells at E12.5. Pparγ, a master regulator of adipogenesis, was reported to be expressed in Myf5 lineage as early as E14 (Wang et al., 2014), while *Pparγ* was not detected by either real-time PCR or RNA-seq at E14.5 in Pax7 lineage. These observations indicate that compared to Myf5 lineage, Contribution of brown adipocytes from Pax7 lineage initiates at a later time point. Importantly, Klf2, a well-known repressor of adipogenic differentiation but not lineage specification by directly repressing *Pparγ2* (Banerjee et al., 2003; Lee and Ge, 2014; Wu et al., 2005), was specifically expressed in Ebf2 group at E14.5, suggesting active mechanisms exist to inhibit early terminal fat differentiation.

The current challenge on investigating lineage-of-interest comes from the lack of robust surface markers and protocols that could isolate and enrich cells purely from that lineage. Pdgfra is regarded as a cell surface marker for brown adipocyte progenitors (Lee et al., 2012), yet we here show that *Pdgfra* is in fact upregulated in the fibroblast-like, dermal, and adipogenic lineages, and the encoded receptor protein can be used as cell surface marker to label all these linages at specific embryonic time points. Pdgfra therefore is not the optimal choice of surface marker to investigate solely adipogenic lineage rising from Pax7 progenitors; however, we found that Thy1 is excellent for enriching cells of the dermal lineage and could be used with Pdgfra for double sorting to separate cells from dermal and adipogenic lineages in E14.5 embryos in Pax7 lineage tracing studies. In line with our cell type validation with real-time PCR studies and *in vitro* cell culture, our findings show it is possible to isolate lineage-of-interest, especially dermal and adipogenic lineages that was previously challenging to isolate, with combination of these two surface markers.

Previously it has been described that cell fate during mouse embryo development could be regulated with transcription factors. MyoD and Myf5 knockdown have been demonstrated to promote cell fate change in muscle progenitors to brown adipocytes (An et al., 2017), suggesting the possibility of cell fate reprogramming in Pax7 lineage at different developmental branch points. We have investigated other potential transcription factors, including Nfatc4 and Pitx3 at E12.5 between myogenic and fibroblast lineages, and Bcl11b and Meo2 at E14.5 between dermal and adipogenic lienages, that could be crucial in cell fate decision especially at different developmental branch point.

Canonical Wnt signaling pathway is crucial for both dermis and epidermis during embryogenesis. Dermal fibroblast at early developmental stage transduces Wnt signal to epidermis and triggers skin appendages development (Andl et al., 2002; Lim and Nusse, 2013; Veltri et al., 2018). Our analysis focusing on dermal cell population in Pax7 lineage is consistent with previous studies which outlines canonical Wnt signaling pathway activities in early dermal fibroblasts and skin development. Disruption of Wnt/β-catenin activity demonstrated to alter dermal layer morphology and hair follicle growth (Andl et al., 2002; Chen et al., 2012). Although the significance of Wnt signaling pathway in dermal development is well-studied, in light of our findings that dermal and adipogenic lineages come from an early fibroblast-like population, we postulate that upregulation of Wnt signaling pathway defines cells from fibroblast-like population as dermal lineage, while the cells have little to no Wnt/β-catenin activity are defined as adipogenic lineage.

In conclusion, we herein present a single-cell transcriptome analysis on Pax7 lineage. Transcriptomic profile of Pax7 lineage outlines cell fate commitment time points of myogenic, dermal, and adipogenic lineages. Differential gene expression analysis supported by gene expression analysis reveals lineage-specific surface markers for future experiments, as well as transcription factors that could determine or reprogram cell fate. Potentially, Wnt signaling pathway could be involved in defining dermal and adipogenic lineages from early fibroblast-like cells. Our discovery promotes further lineage-specific investigations of Pax7 progenitor cells.

## Materials and methods

### Animals

Pax7^creER^ (Gaka) (stock 017763) and R26-stop-EYFP (stock 006148) mice were from Jackson Laboratory (Bar Harbor, ME, USA). Mice were housed in the Animal and Plant Care Facility (APCF) at HKUST. All the experimental protocols were approved by APCF. A single dose of TMX (75μg/g body weight) were injected intraperitoneally into the pregnant mice at E9.5 to label Pax7 expressing cells. Embryos were harvested at E10.5, E12.5 and E14.5.

### Embryonic cell isolation and flow cytometry

Isolated mouse embryos were minced and digested in sorting medium (Ham’s F10 with 10% horse serum) containing 400U/ml Collagenase II (Worthington; LS004177) for 1h at 37 °C in a shaking water bath. For E14.5 or older embryos, additional Dispase (1U/ml; Gibco) was added. The digested embryos were then washed in the sorting medium and filtered through a 40 μm cell strainer (BD Falcon). After centrifugation, the cells were suspended in sorting medium and stained with antibodies for 45 min. The cells were then sorted on a BD FACSAria III or Influx™ cell sorter. Antibodies used for staining: Pdgfrα-PE 1:200 (BioLegend; 135905), Thy1.2-APC 1:200 (BioLegend; 140311), Itga7-APC 1:300 (Ablab; 67-0010-10).

### Library preparation and sequencing

Both Smart-seq2 (Picelli et al., 2013) and 10X library preparation protocol (10X Genomics) were used to construct single-cell libraries of Pax7-lineage cells. For Smart-seq2, YFP+ cells were sorted into 96-well plates and proceeded immediately to reverse transcription and cDNA amplification as previously described; however, we observed excessive amount of primer-dimer post-amplification, hence every primer volume was reduced by 10%. Libraries were then completed with Illumina Nextera XT library preparation kit (Illumina) and sequenced in Nextseq 500 (Illumina) at average of 2.69 million reads per cell. For 10X, cells were sorted into single-cell suspension and proceeded immediately to library preparation with 10X 3’ kit and 10X chromium controller (10X Genomics). Libraries were sequenced in Nextseq 500 (Illumina) at average of 74 thousand reads per cell.

### Data processing

Smart-seq2 libraries were demultiplexed according to index used to generate FASTQ files. FASTQ files were quantified to transcript level expression using Kallisto (Bray et al., 2016). then summarized to gene level expression matrix with R package *tximport* (Soneson et al., 2016). Cells without YFP expression were discarded, resulting total of 728 cells. 10X libraries were demultiplexed and counted for genes using Cell Ranger (10X Genomics) resulting total of 3140 cells. Both sets of data were processed and analyzed using *Seurat* package (Butler et al., 2018). Log-normalization and linear regression were performed on both datasets. Smart-seq2 data were regressed with sequencing reads and cell-cycle score, while 10X data were regressed with number of UMIs per cell and cell-cycle score. Cell-cycle was scored on strategy previously described implemented in *Seurat* package (Tirosh et al., 2016). After processing data, we first selected variable genes using a dispersion threshold Log variance-mean ratio to perform principal component analysis (PCA) to project data into low dimensional subspace, and selected numbers of principal components that explained high variance. The low dimensional subspace was then used to calculate Euclidean distance metric which was used to visualize cells by t-distributed stochastic neighbor embedding (t-SNE) and cluster cells by SNN clustering.

### Gene Ontology enrichment analysis

Gene Ontology enrichment analysis were performed using PANTHER (Mi et al., 2017). Gene Ontology terms were annotated by Gene Ontology Consortium (Ashburner et al., 2000; The Gene Ontology Consortium, 2017). GO terms with adjusted p-value less than 0.01 were reported.

### Differential expression analysis

Differentially expressed genes were identified with likelihood-ratio test for single-cell experiments previously described (McDavid et al., 2013). Genes with adjusted p-value less than 0.01 were reported. Population of test was based on number of cells in cluster.

### Cell culture

For embryonic cell culture, the 48-well plates were pre-coated with 0.1% at 37°C for 1h. The sorted embryonic cells were cultured in Dulbecco’s modified Eagle’s medium (DMEM) with 10% fetal bovine serum (FBS). The adipogenic differentiation medium (ADM) was described previously (An et al., 2017). After reaching confluency, the cells were first cultured in ADM I for 2 days and then switched to ADM II for 2 days or more.

### Oil Red O staining and immunostaining

For Oil Red O staining, cells were fixed in 10% formalin, rinsed with ddH_2_O and stained in Oil red O working solution (final 36% in Triethyl phosphate). The cells were then incubated with haematoxylin for 5 mins for nuclei staining. For immunostaining, cells were fixed with 4% paraformaldehyde for 5 mins, followed by permeabilization in 0.5% PBST and blocking in 4% BSA. Primary antibodies used are: Myod1 1:200 (Dako; M3512), Ebf2 1:100 (R&D; AF7006)

### RNA isolation and RT-qPCR

Total RNA from embryonic cells were extracted with TRIzol method, followed by cDNA synthesis using ImProm-II reverse transcription system. Real time PCR were performed on a Roche LightCycler^®^ 480 machine using SYBR green master mix.

## Acknowledgements

We thank We thank Prof. Can Yang from Department of Mathematics for his input in data analysis, and lab member Yitai An and Xiuqing Wei for their aid in sorting, as well as all other members for their helpful discussion and generous support.

