## Supplementary figures for "Cell Fate Determining Molecular Switches and Signaling Pathway in Pax7-expressing Somitic Mesoderm"

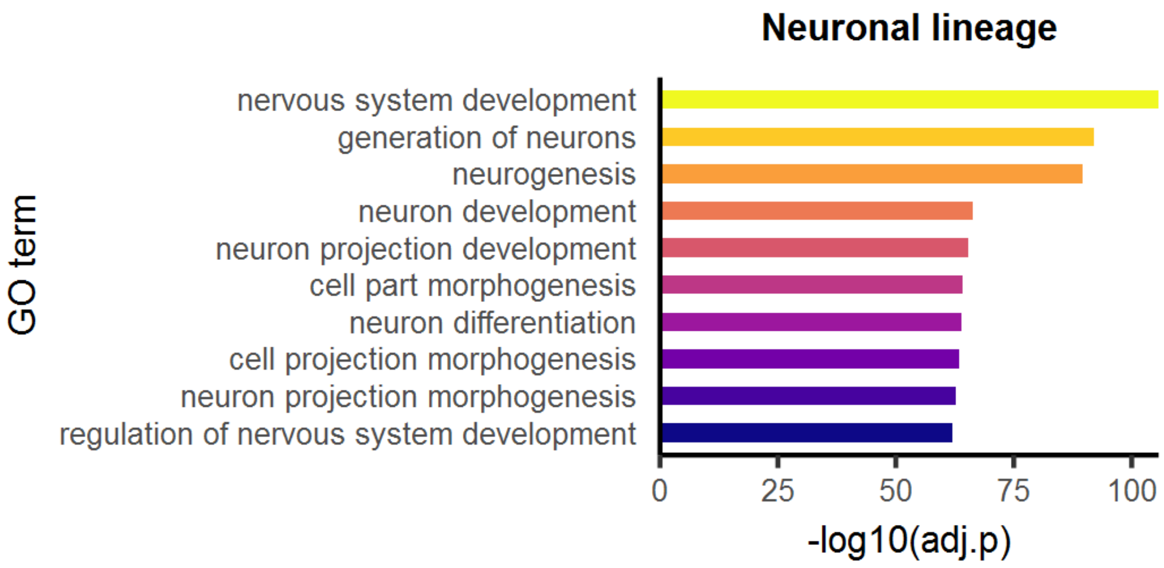


**Fig. S1. Gene Ontology enrichment analysis on neuronal lineage.** Top 10 terms with lowest p-value were related to neuron development.

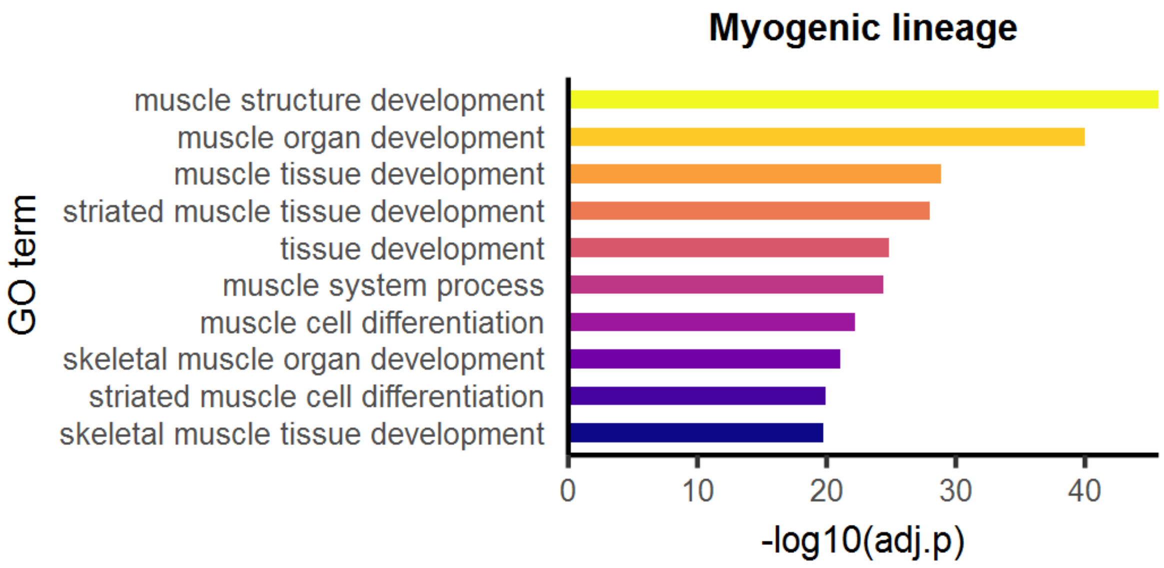


**Fig. S2. Gene Ontology enrichment analysis on myogenic lineage.** Top 10 terms with lowest p-value were related to muscle structure development, and muscle cell differentiation.

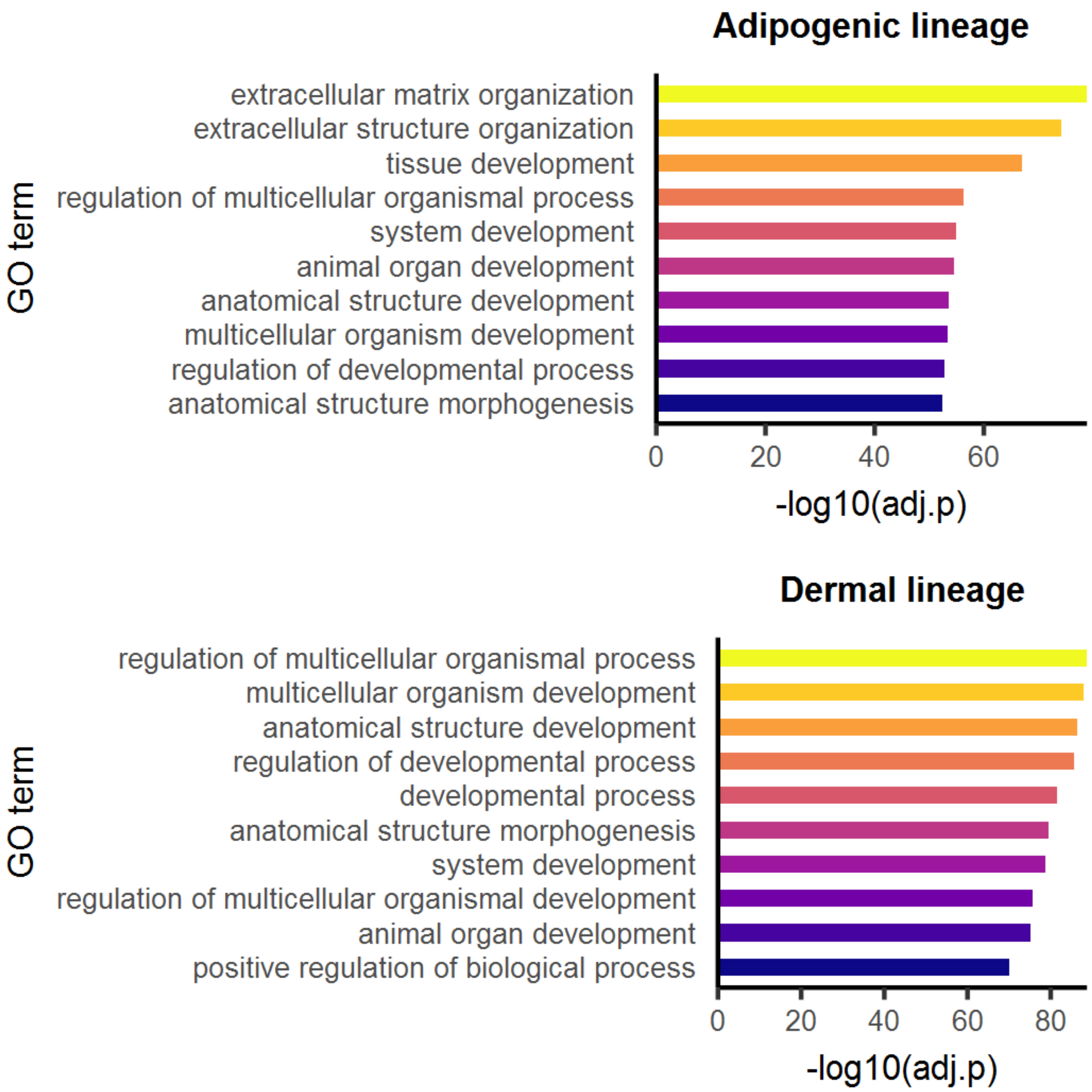


**Fig. S3. Gene Ontology enrichment analysis on adipogenic and dermal lineages.** Top 10 terms with lowest p-value were not related to corresponding lineage or cell types, with terms not specific to certain tissue type.
